# Morphological plasticity of endophytic *Chitinophaga pinensis*

**DOI:** 10.1101/2025.07.09.663833

**Authors:** Janine Liedtke, Frans Rodenburg, Chao Du, Le Zhang, Gilles P. van Wezel, Ariane Briegel

## Abstract

Environmental changes, whether due to climate change or human influences, compromise the resilience of plants to biotic and abiotic stresses, such as pathogens, drought and heat. Plant microbiota are known to promote plant resilience. To be able to harness the power of the plant microbiome we need to identify microbiota with health-promoting properties. Recent studies have demonstrated that the bacterium *Chitinophaga pinensis* enhances plant health and increases resistance to fungal infections. Here, we show that *C. pinensis* exhibits an unusually high morphological plasticity, switching between a filamentous and a spherical cell state, each of which is characterized by a distinct transcriptional profile. Despite these transcriptional differences, spherical cells remained metabolically active and replicating, while lacking structural characteristics typically associated with dormant states. Furthermore, the spherical cell morphology of *C. pinensis* facilitates hitchhiking behaviour and motility via surfactin cheating, potentially influencing its dispersal and interactions within the plant microbiome. To investigate the structural dynamics and transcriptional adaptation of this plant endophyte, we applied a combination of microscopy and culture-based techniques. Taken together, our study provides new insights into the morphological flexibility and transcriptional regulation of the plant-beneficial *C. pinensis*.

## Introduction

Modern agriculture increasingly faces challenges due to climate change and a growing world population. Therefore, new measures and innovative strategies must be developed to conserve resources and minimize losses to ensure a sustainable future.

One of these strategies is to understand and utilize the natural defence and resistance mechanisms of plants. Numerous studies have already shown that the plant microbiota plays a decisive role in this context (reviewed in ref. [1–3]). Plants and their associated microbiota are often referred to as a holobiont, as they engage in complex symbiotic interactions and have an evolutionary history as a community (reviewed in ref. [4]). Among the most abundant phyla within the plant microbiota, the phylum Bacteroidetes stands out as it contains numerous symbiotic strains associated with various hosts [5, 6].These microorganisms can enhance host fitness either directly by promoting plant health or indirectly by supporting other members of the microbiota. Those that colonize plant tissue without causing harm or negatively affecting the host are known as endophytes. Recent studies have shown that the endophytic strains *Flavobacterium anhuiense* and *Chitinophaga pinensis* contribute to plant health and improve stress tolerance [7]. The diversity of the plant-associated microbiota reflects a range of strategies that can enhance plant resilience. One such strategy is the “cry-for-help” response, in which plants actively recruit beneficial microbes by releasing root exudates in response to environmental stress. The composition of these exudates varies depending on the stress conditions, which leads to a specific metabolic adaptation of the bacteria already residing in the microbiota and stimulates the production of secondary metabolites [1, 8–10]. However, effectiveness of induced stress tolerance depends on the microbiota composition, and missing key members can decrease the response effect [11]. This effect is even more pronounced during activated defence mechanisms in the presence of plant pathogens [10, 12]. In order to optimize the composition of the microbiota to protect plants naturally, it is important to identify key microbiome members and understand the factors involved in their recruitment.

As mentioned above, plants excrete root exudates to actively recruit and shape their microbiota according to their needs [8]. These exudates contain diverse compounds, including sugars, amino acids, enzymes, fatty acids and polyphenols such as flavonoids [13]. To successfully colonize the rhizo- and endosphere, microbes must be able to detect and respond to those signals. Many motile bacteria achieve this through chemotaxis, allowing them to sense and move along chemical gradients via chemosensory pathways that regulate flagellar motors [14, 15]. Depending on their structural adaptations, bacteria utilize different motility strategies, including flagella-driven swimming and swarming or gliding motility, which relies on surface adhesins [16, 17]. However, some bacteria do not possess chemotaxis arrays or motility mechanisms. This raises the question of how these strains reach the plant and establish themselves within the microbiome. Previous research demonstrated that non-motile spores of endophytic *Streptomyces* can be transported by motile bacteria, a process known as hitchhiking [18]. Hitchhiking allows non-motile bacteria to benefit from chemotactic movements without expending their own energy, potentially providing ecological advantages in host colonization. As proposed by Muok *et al*. (2021) [18] and Seymour (2024) [19], hitchhiking can also yield in long-term benefits for bacterial communities, facilitating movements through barriers and promoting metabolic exchange.

Here, we focus on *Chitinophaga pinensis*, which was recently identified as an important member of the plant microbiota particularly regarding pathogen resistance and stress tolerance [7]. Originally isolated from pine litter, *C. pinensis* is a Gram-negative bacterium found in the endosphere of various plants, including crops [7]. It is known for its ability to degrade complex polysaccharides such as chitin and produce a variety of biologically active molecules, highlighting its potential value for agriculture and industrial applications [20]. Furthermore, previous studies have reported that *C. pinensis* forms so-called myxospores or microcysts, which have been hypothesized to represent a resting stage similar to bacterial spores [21–23]. However, these structures have primarily been characterized based on their spherical morphology under the light microscope, and their metabolic state remained unclear. While most research on *C. pinensis* focused on its metabolic capabilities alone, in this study we investigated the ultrastructural morphology of cells as well as their interactions within the holobiont using a combination of microbiological assays, cryo-electron microscopy and transcriptomics. In particular, we aimed to gain insight into whether the morphology of *C. pinensis* depends on the growth conditions, and see how this correlates to the straińs ability to interact with plant tissue. Additionally, we aimed to determine whether these interactions are solely based on chemical signals or if physical structures, such as outer membrane vesicles [24], play a role in the communication with the plant. Notably, the *C. pinensis* strain studied by Carrión (2019) [7] is neither motile nor equipped with chemotaxis arrays, which raises questions about its recruitment and ecological role within the holobiont. Finally, we investigated how these characteristics of *C. pinensis* contribute to its ecological function and potential impact on the dynamics of the plant microbiome.

## Results

### *Chitinophaga pinensis* shows two distinct cell morphologies

To gain a better understanding of the general morphology of *C. pinensis* over time, we investigated the cell culture at different incubation times by light microscopy. In fresh cultures up to 20 h of growth, the cells are filamentous and can reach lengths of up to 40 µm, as described previously [21]. Intriguingly, between 20 and 40 h of growth, the morphology of *C. pinensis* dramatically transformed from several µm long filamentous cells to small spherical cells with a diameter of 600–700 nm (Figure 1). When transferred to fresh media with a lower cell density, the spherical cells reverted to the filamentous cell state (Figure S1). To gain a more detailed insight into these two cell morphologies, we used cryo-electron tomography (cryo-ET). Both cell types were enclosed by a typical Gram-negative cell envelope, enclosing a typical cytoplasm filled with evenly distributed ribosomes and other cellular content. Neither cell type resembled previously described spores or cysts, such as for example the presence of a thicker protective cell envelope (Figure 1).

**Figure 1.**
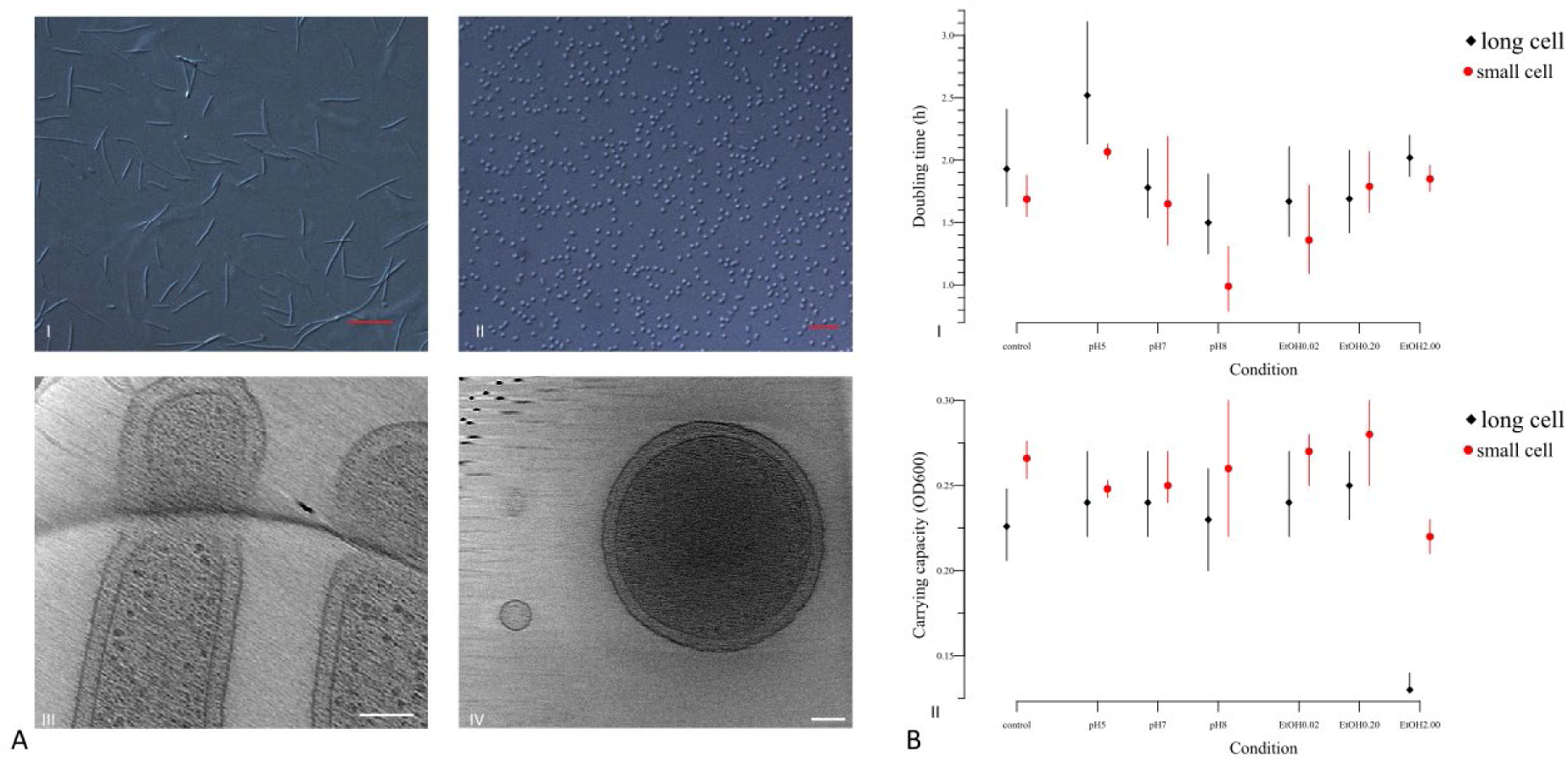
Morphological plasticity and stress resistance for the *C. pinensis* morphotypes. Cells were imaged by light microscopy (I – AII) and cryo-electron tomography (AIII – AIV; scale bar 100 nm). (AI) Long filamentous cells after 20 h (scale bar 10 µm) and small spherical cell shape after 40 h of incubation (scale bar 5 µm). Subsequently differences in resistance behaviour were tested in both morphologies in 0.1x TSB medium supplemented with increasing concentration of ethanol or change of media pH. Measured were the change of OD_600_ over 20 h and displayed are the calculated growth parameters – doubling time (BI) and carrying capacity (BII) based on fitted growth curves with a confidence interval of 95%. (N= 3)

To better visualize the morphological switching of *C. pinensis* from filamentous to spherical growth, we aimed to introduce GFP into the bacterium. The GFP labelling was designed to enable live-cell imaging and precise tracking of morphological changes, particularly since the spherical cells are very small (600–700 nm), making them difficult to keep in focus and distinguish from other particles or debris under the microscope. Therefore, the gene encoding GFPmut3, under the control of its native *gap* promoter, was cloned into pCP23 (details in Materials and Methods), a shuttle vector between *Flavobacteria* and *E. coli*. The resulting construct, pGWS1802, was successfully introduced via electroporation. However, the plasmid was not stably maintained: green fluorescence diminished with each subsequent bacterial generation. This suggests that the plasmid failed to replicate in *C. pinensis*, most likely due to incompatibility of the repB replication origin on pCP23, which is optimized for Flavobacteria and does not function effectively in this species.

### Resistance profiles of cell morphologies to environmental and chemical stressors

The small spherical cells observed after prolonged growth were consistent with previous reports, which described them as microcysts or spores. However, our cryo-ET results revealed that the small spherical cells forms do not have a typical spore morphology, such as a thickened cell envelope and tightly packed cytoplasm content. Therefore, we wondered whether the two cell morphologies exhibit distinct responses to various physical and chemical stress factors. Both cell morphologies exhibited similar resilience to the following stress factors: UV light irradiation for 10 minutes, heat stress at 60 °C and sonication for up to 5 min. Both cell morphologies of *C. pinensis* survived after desiccation and could be revived by plating on fresh medium. Light microscopic examination of the desiccated *C. pinensis* cells revealed small spherical cells, independent of their initial morphology. Further chemical stress tests showed slight differences in resistance behaviour of the two cell morphologies in response to changes in the pH and in the presence of up to 0.2% ethanol (Figure 1). However, in the presence of 2% ethanol both cell morphologies grew slower, with the OD_600_ ≤ 0.25. Neither cell morphology of *C. pinensis* grew in SDS-containing (0.02–2%) media.

Overall, the spherical morphology exhibited higher carrying capacity and a slightly shorter doubling time compared to the filamentous cell morphology (Figure 1). Nevertheless, the differences in the resistance profiles between the two morphologies were marginal.

Next, we investigated whether the decrease in cell size and the formation of small spherical cells represented a stress reaction. For this purpose, growth in the presence of salt (0.5–2%) or chloramphenicol (12.5–100 µg/ml) was analysed. When cells with a filamentous phenotype were exposed to higher salinity and chloramphenicol concentrations, the cell size decreased (Figure S3). With increasing salt and chloramphenicol concentrations, the spherical cells tended to dominate the culture faster compared to the control conditions (Figure S3). Regardless of the salt and chloramphenicol concentration in the medium, spherical cells appeared in all cultures after 2 days of incubation.

### Transcriptomic analysis of 20 h and 40 h cultures

Spherical cells were previously assumed to be dormant spores. However, the cryo-EM and resistance assays presented here revealed no structural or physiological features typically associated with dormancy. To assess the changes in transcriptional patterns between the two time points and corresponding culture types, we performed transcriptomic analysis on RNA obtained from cultures harvested after 20 h and 40 h of growth, dominated by filamentous and spherical cells, respectively. All cultures were grown under identical conditions (1.0x TSB, 25 °C, 250 rpm) (Figure S1). It is important to note that the growth rate of the cultures was comparable.

Principal component analysis (PCA) showed a clear transcriptional distinction between the two time points (Figure S4). Differentially expressed genes (DEGs) were defined as showing a fold change of ≥ 2 (FC ≥ 2) and an adjusted *p*-value (padj) of 0.05. Based on these criteria, 691 genes were upregulated and 615 downregulated in the 40-h samples compared to the 20-h samples of a total of 6094 genes (Figure S5). The lists of up-, down-regulated DEGs and all DEGs were subjected to KEGG pathway enrichment analysis (Table 1). Genes downregulated at 40-h were significantly enriched for ribosomal components (cpi03010), and upregulated genes were depleted for those involved in amino acid biosynthesis (cpi01230). Additionally, genes involved in fructose and mannose metabolism (cpi00051) and fatty acid biosynthesis (cpi00061) were predominantly downregulated. Within the fatty acid biosynthesis pathway, most of the identified genes (19 out of 25) showed reduced expression, including 9 DEGs, indicating a broad transcriptional shift in this pathway.

**Table 1.**
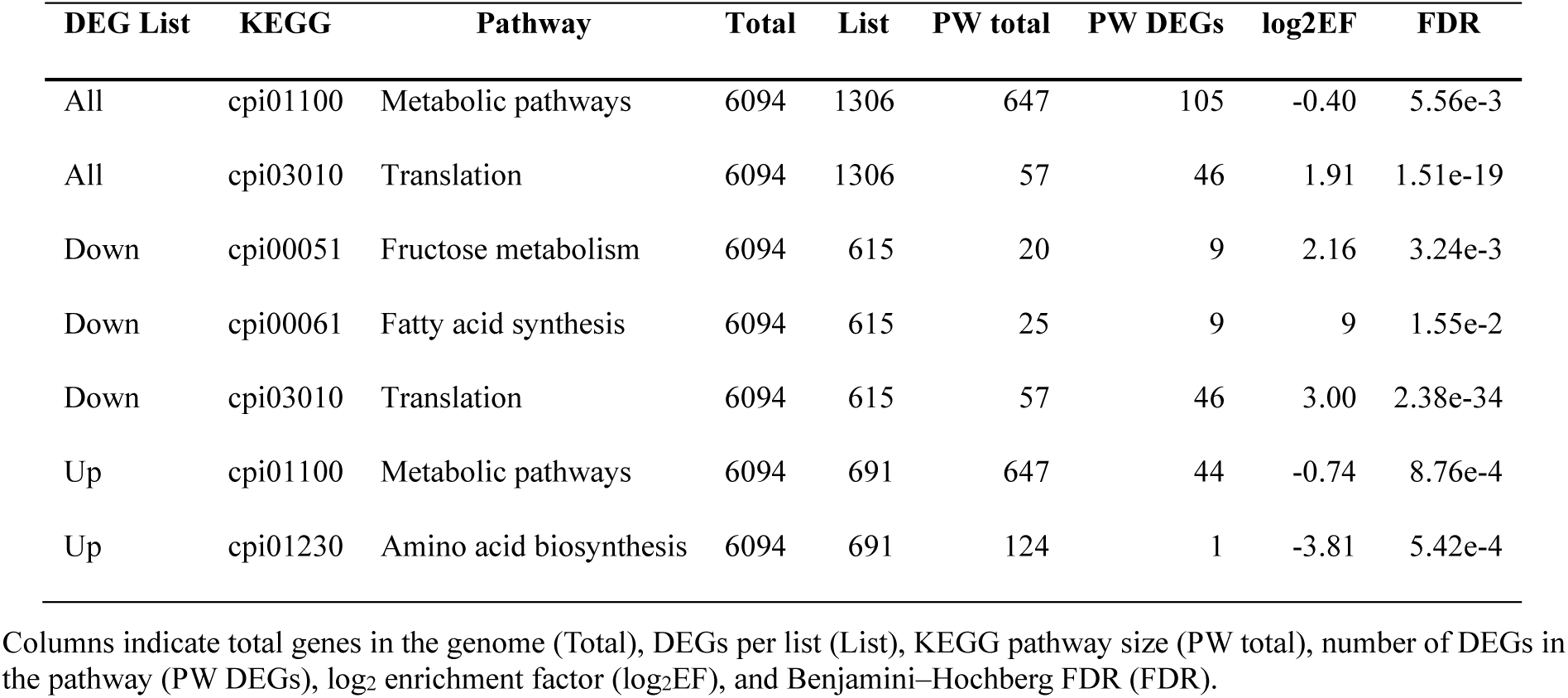
KEGG pathway enrichment analysis of differentially expressed genes (DEGs).

| DEG List | KEGG | Pathway | Total | List | PW total | PW DEGs | log<sub>2</sub>EF | FDR |
| --- | --- | --- | --- | --- | --- | --- | --- | --- |
| All | cpi01100 | Metabolic pathways | 6094 | 1306 | 647 | 105 | -0.40 | 5.56e-3 |
| All | cpi03010 | Translation | 6094 | 1306 | 57 | 46 | 1.91 | 1.51e-19 |
| Down | cpi00051 | Fructose metabolism | 6094 | 615 | 20 | 9 | 2.16 | 3.24e-3 |
| Down | cpi00061 | Fatty acid synthesis | 6094 | 615 | 25 | 9 | 9 | 1.55e-2 |
| Down | cpi03010 | Translation | 6094 | 615 | 57 | 46 | 3.00 | 2.38e-34 |
| Up | cpi01100 | Metabolic pathways | 6094 | 691 | 647 | 44 | -0.74 | 8.76e-4 |
| Up | cpi01230 | Amino acid biosynthesis | 6094 | 691 | 124 | 1 | -3.81 | 5.42e-4 |

Columns indicate total genes in the genome (Total), DEGs per list (List), KEGG pathway size (PW total), number of DEGs in the pathway (PW DEGs), log<sub>2</sub> enrichment factor (log<sub>2</sub>EF), and Benjamini–Hochberg FDR (FDR).

Despite the observed downregulation of genes involved in translation and biosynthesis processes at 40-h, growth measurements under control conditions showed similar growth rates for both time points. The doubling time was 1.53 h at 20-h (µ = 0.45 h^-1^, 95% CI: 1.32 – 1.83), and 1.62 h at 40-h (µ = 0.43 h^-1^, 95% CI: 1.39 – 1.94). The confidence intervals overlapped.

### Morphological variation under different culture conditions

Next, we sought to determine the factors that trigger morphological changes. For this, we systematically varied (i) cell density (OD_600_ = 0.2 and 0.5), (ii) nutrient content, and (iii) the presence of quorum sensing (QS) signals using five different media (FM, LM, SM, SMT, B). As these factors are interrelated, they were tested by adjusting their concentrations in the media.

The general pattern was a decrease in cell length of approximately 29% for filamentous cells and an increase of about 74% for spherical cells over the measured time (Figure 2).

**Figure 2.**
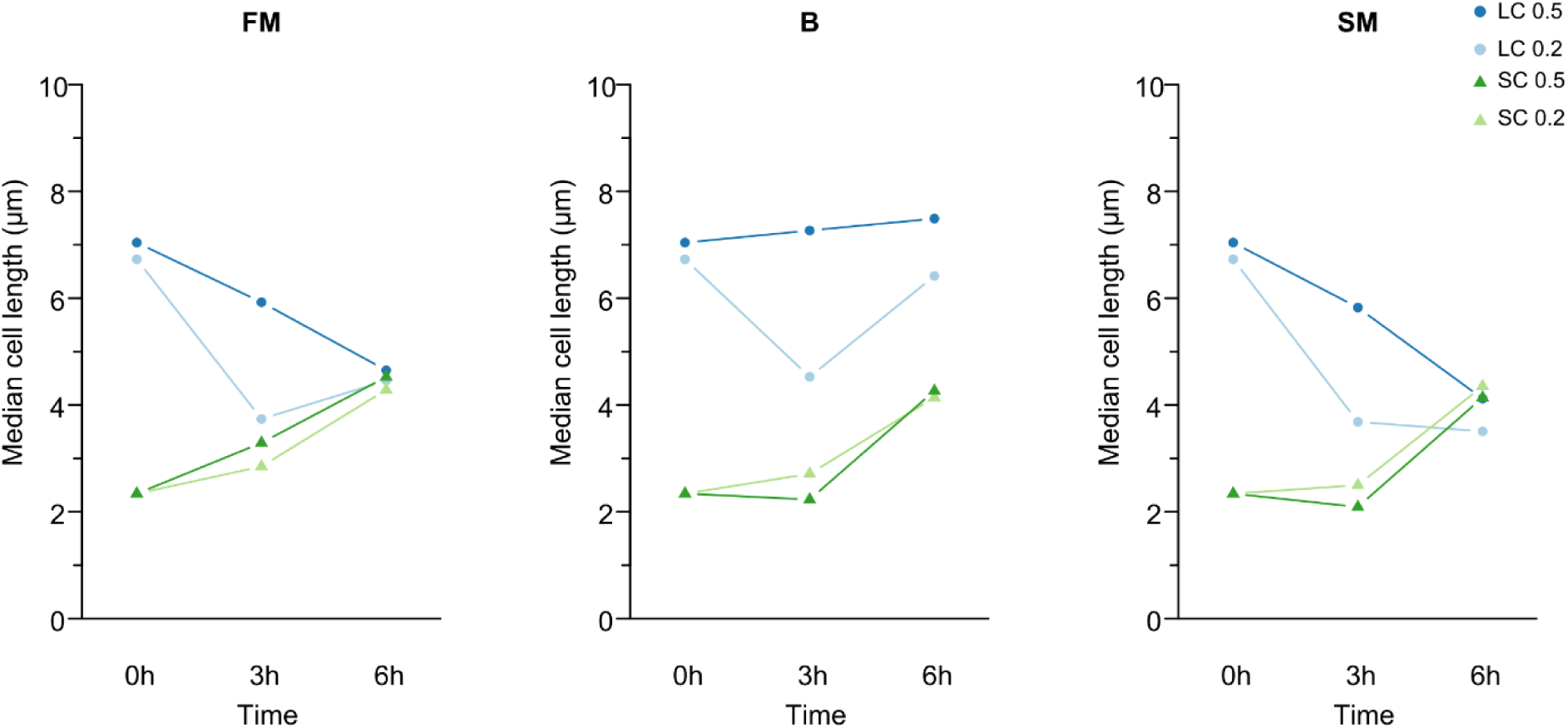
Trigger test of morphological change. Observed pattern of change in cell morphology under different growth conditions (N = 3). Shown is the determined median of cell length for each cell morphology at the cell densities of OD_600_ = 0.2 and 0.5 at the measured time points: 0, 3, and 6 h. (A) Illustration of the predominant pattern of cell length change observed in the different media, here using FM as an example. Deviating pattern were observed in the media B and SM for filamentous cells. (B) Illustration of the pattern of filamentous cells with an initial OD_600_ = 0.5 in medium B. (C) Deviating pattern of cell length change from the general pattern of filamentous cells at a cell density of OD_600_ = 0.2 in medium SM. The median cell length is given in µm.

For both morphologies, measurable changes in cell length were observed after 3 h compared to the initial values, and density-related differences became apparent. After a total of 6 h, these differences decreased, and the cell lengths showed a trend towards convergence within each cell morphology (Figure 2).

The influence of cell density was more pronounced in filamentous cells than in spherical cells. In spherical cells, both densities led to a similar increase in length, differing primarily in the speed of increase. In contrast, filamentous cells showed different patterns depending on the initial density; at high density (0.5), cell length decreased continuously by approx. 32% after 6 h, while at low density of (0.2), the cell length initially decreased by 45% after 3 h, followed by an of approx. 27% after 6 h. Deviations from this general pattern were observed in B and SM media, whereby in medium B at higher cell density (0.5), filamentous cells increased in length by approximately 6% over time (Figure 2B), while in SM media at lower cell density (0.2), filamentous cells continued to decrease in length after 6 h (approx. 48%) (Figure 2C).

During the experiment, we assumed that the nutrient content of the media decreased in the following order, from the highest to the lowest content: SMT > FM > LM > SM > B. Whereby medium B was assumed to contain neither nutrients nor QS signals. We also presumed that LM contained QS signals from filamentous cells, while SM and SMT contained QS signals from spherical cells.

Overall, changes in cell length were observed in both cell morphologies under all conditions tested. Comprehensive details and individual diagrams for each condition are provided in the supplementary materials Figure S6.

### *C. pinensis* spherical cells are dispersed by *B. subtilis*

Based on our observation that *C. pinensis* is non-motile under the condition tested and exhibits morphological plasticity, we next investigated whether the spherical cells could provide an adaptive advantage, particularly in terms of their potential to hitchhike on motile bacteria.

Here, we repeated the hitchhiking assay according to Muok *et al*. (2021) [18] in order to determine the hitchhiking ability of the two cell morphologies of *C. pinensis*. For this purpose, filamentous or spherical cells of *C. pinensis* were isolated and placed on semi-solid media plate over *B. subtilis*. The hitchhiking behaviour of *C. pinensis* cells was assessed by observing their spreading indicated by the yellow-orange colonies.

As shown in Figure 2, when spherical cells were added, the yellow colonies spread across the plate. When filamentous cells of *C. pinensis* were plated, the colonies remained in the centre of the plate.

To understand the nature of the hitchhiking behaviour, we tested the ability of *C. pinensis* to hitchhike in the presence of *B. subtilis* mutants that were impeded in their movements either by inhibition of surfactin production (Δ*espH srfAA*::mls), suppression of flagellar motility (Δ*motAB*}) or absence of flagella (Δ*hag*}). In addition, a laboratory strain of *B. subtilis* (DK605) that had lost its swarming ability [18] was included in this study. The test revealed that the hitchhiking behaviour of *C. pinensis* is independent of flagellar movement and the presence of flagella. While the spherical cells can be dispersed by swarming and sliding motility of *B. subtilis*, they cannot be dispersed by a *B. subtilis* mutant (Δ*espH srfAA*::mls) that is limited to swimming motility and unable to produce surfactin. A surfactin cheating assay demonstrated that the spherical cells of *C. pinensis* follow surfactin dispersal in and on media surface (Figure 3).

**Figure 3.**
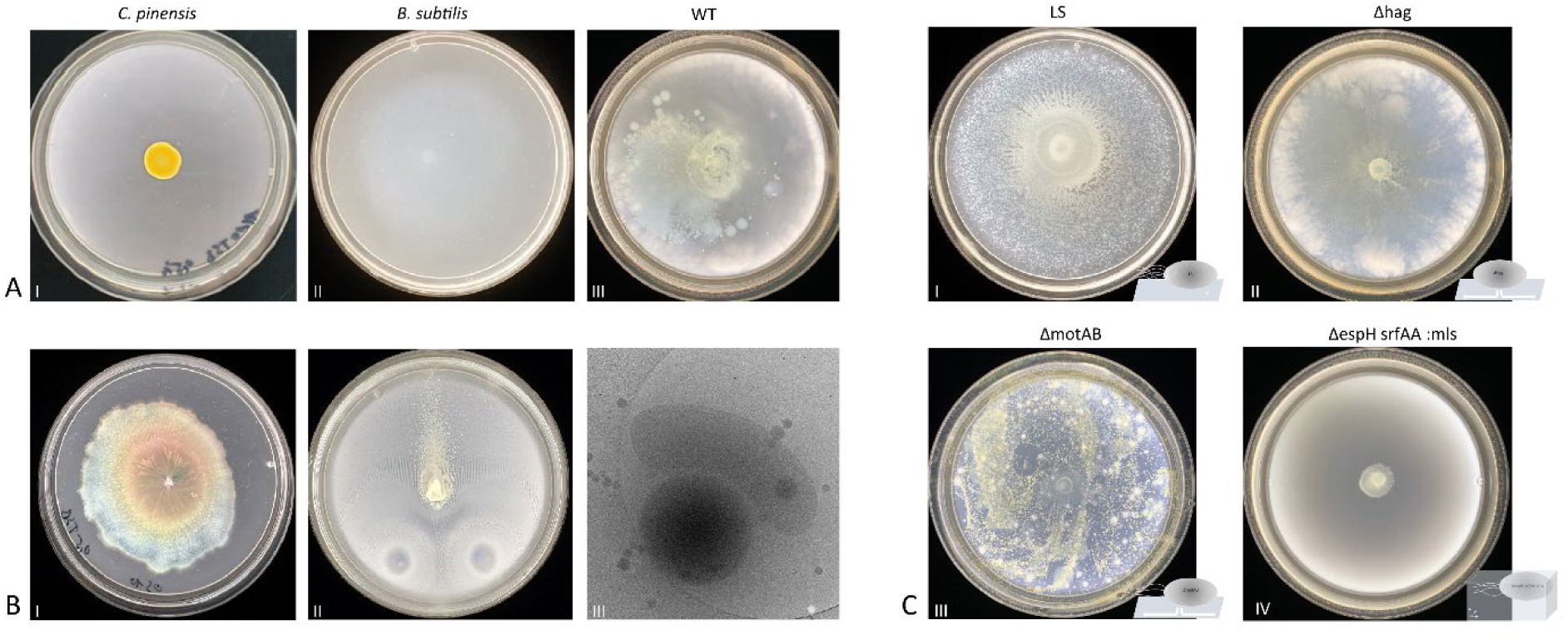
Motility assay showing hitchhiking behaviour and passive translocation. *C. pinensis* strain (AI) did not display motility under the tested conditions and remained at the point of inoculation. But when combined with a motile bacteria like a wild type (WT) *B. subtilis* (NCIB3610) (AII), dispersal of *C. pinensis* appeared over the plate (AIII). Further hitchhiking tests were conducted with *B. subtilis* mutants, that were either hindered in their motility by either inhibition of surfactin (*espH srfAA*\::mls) (CIV), suppression of flagella motility (*motAB*) (CIII) or absence of flagella (*hag*) (CII). Additionally, a lab strain (LS; DK605) that lost its swarming capability was included (CI). The test revealed flagella-independent hitchhiking behaviour of *C. pinensis*. However, this behaviour could not be observed in combination with a swimming-only strain as carrier (CIV). Subsequently, the surfactin-cheating capability of *C. pinensis* was tested (BI) and this demonstrated that it follows the crowd movement of *B. subtilis* (NCIB3610) (BII). Furthermore, the hitchhiking behaviour of *C. pinensis* with *B. subtilis* (DK605) were imaged by cryo-electron microscopy (BIII).

Next, we tested whether the spherical cells of *C. pinensis* follow the surfactin spreading in the presence of *B. subtilis* cells, assuming that the surfactin produced by *B. subtilis* cells is distributed over the entire plate due to their movement. Despite the presence of surfactant on the entire plate, *C. pinensis* cells only spread along the movement of *B. subtilis* (Figure 3). When the spherical cells were placed between two *B. subtilis* inoculation sites, the *C. pinensis* colonies remained between them. Although the spherical cells of *C. pinensis* can cheat using *B. subtilis* surfactin, they cannot spread in the opposite direction of *B. subtilis* movement (Figure 3).

## Discussion

### *C. pinensis* has two morphologies with distinct transcriptional profiles

*C. pinensis* is described in the literature as a long filamentous bacterium that can produce small spherical bodies, often referred to as myxospores, microcysts or spores, and assumed to represent a dormant stage [21–23]. Although these terms are frequently used, there is neither a clear definition nor a distinction between them [25]. Dormancy is defined as an adaptive state of reduced metabolic activity and arrested growth that enhances survival under unfavourable conditions [26]. It is typically associated with structural and physiological adaptations such as thickened cell walls, condensed DNA, and enhanced resistance to environmental stressors [26]. However, stress resistance assays and structural analysis suggest that morphological changes in *C. pinensis* do not follow the classical characteristics of dormancy-associated differentiation. Our transcriptomic analysis indicates a shift in gene expression patterns in spherical cells, suggesting a metabolic slow down, although cells continue to proliferate and maintain distinct transcriptional activity. Furthermore, there was no major difference in growth rate between 20 h and 40 h cultures, dominated by filamentous and spherical cells, respectively. These results suggest that *C. pinensis* does not really enter a dormant phase but instead modulates its transcriptional profile at the molecular level as part of an adaptive response to changing nutrient availability and environmental conditions.

Morphological plasticity is a well-documented survival strategy among bacteria, enabling them to respond dynamically to environmental change. Some bacteria undergo sporulation under unfavourable condition, whereas others alter their morphological characteristics to optimize resource acquisition, evade predation, or adapt to fluctuating nutrient availability. *Escherichia coli*, for example, can undergo filamentation under stress to avoid predation or shrink size to minimize metabolic demands [27, 28]. In this context, the ability of *C. pinensis* to transition between morphologies may provide similar ecological advantages, contributing to its persistence in diverse environments. Beyond individual survival, morphological transitions can also play a role in microbial community dynamics and interactions with host organisms. Previous studies have shown that microbial morphology can influence biofilm formation, resource competition, and interspecies interactions within complex microbiomes [29–31]. However, as noted by Shah (2019) [32], the ecological consequences of morphological differentiation in bacteria remain underexplored. In the case of *C. pinensis*, the impact of its morphological states on its functional role within the plant microbiome remains unclear.

### Relevance of morphological and transcriptional plasticity for host*–*microbiota interactions

Previous studies on morphological plasticity have primarily focused on stress responses, often overlooking concurrent metabolic changes [31–33]. Consequently, the link between morphology and transcriptional changes beyond stress adaptation remains insufficiently explored. The observed morphological and transcriptional plasticity in *C. pinensis* challenges conventional assumptions, as filamentation is typically associated with stationary phase adaptation, where secondary metabolite production is prioritized over cell growth and occurs as a stress response [34]. In this way, bacteria ensure that secondary metabolites are produced at low cost and with high benefit [35]. However, in *C. pinensis*, filamentation and increased transcription of secondary metabolites occurs during early growth when nutrients are abundant, and cell density is low. Filamentation is energetically costly and can reduce fitness and carrying capacity [31, 35], suggesting it must provide compensatory benefits in *C. pinensis*. Filamentation has been associated with several ecological advantages, including improved surface interactions, biofilm formation, competitive polymer degradation, nutrient acquisition in heterogeneous environments, and a possible intracellular spreading mechanism [36–38]. The interplay between morphological and transcriptional plasticity in *C. pinensis* reflects a complex regulatory network within the holobiont, likely contributing to microbial stability and adaptation. However, further research is needed to determine whether these morphological transitions provide ecological advantages in competitive environments and how they influence plant-microbe interactions [34, 39].

### Hidden trigger of recurrent morphological change

To identify triggers for morphological change beyond stress factors, three commonly described factors – nutrient availability, QS signalling, and cell density – were tested [30, 40]. Since these factors co-occur, they could not be tested in isolation. Instead, their concentrations were varied, and the speed of morphological change was analysed. Despite these efforts, no clear trigger could be identified, suggesting a complex interplay of growth conditions and regulatory mechanisms. Previous studies have shown that triggers are only effective under certain conditions. If these conditions are not met, the cell’s regulatory mechanisms intervene and prevent the response [31]. This could explain why the filamentous cells in SMT increased in size after six hours, whereas in SM, they continued to decrease (Figure S6). Both media contain QS signals from spherical cells but differed in nutrient composition. Additionally, this effect was absent in filamentous cells at higher initial cell densities, which is a further indicator for a complex interplay of regulatory mechanisms.

Triggers may also be masked by secondary effects. It was suggested that high phosphate concentrations may trigger filamentation of *Paraburkholderia elongata* [31]. However, further analysis revealed that the actual trigger was magnesium deficiency, resulting from intracellular polyphosphate chelation. Similarly, our findings suggest that morphological changes in *C. pinensis* are linked to cell density. This is supported by observations that filamentous cells exhibited density-dependent differences in cell length dynamics, while spherical cells increased in size under all tested conditions (Figure S6). This may be due to dilution effects, as spherical cells were harvested from a high-density culture, potentially initiating morphological transitions.

Growth rate, rather than cell density alone, likely plays a key role in morphological changes as well [34, 38, 40]. The metabolic sensor UDP-glucose has been shown to link carbon availability to growth rate by controlling FtsZ ring assembly, a key regulator of bacterial cell division [34, 38, 40]. However, *C. pinensis* is genetically not readily tractable and despite extensive attempts we could not express FtsZ-GFP.

### Spherical cells can hitchhike and potentially utilize chemotaxis of others

Although motility via the type IX secretion system (T9SS) is known in other *C. pinensis* strains [41], it could not be induced in the strain used in our experiments. This limitation led us to explore whether morphological plasticity provides alternative advantages beyond metabolic efficiency. Given their small size (600–700 nm), the spherical cells of *C. pinensis* could be transported by vascular pressure within plant tissue. However, their movement in soil remains unclear. Without chemotaxis genes, they may rely on other motile bacteria for translocation. Previous studies have shown that some non-motile bacteria exploit the motility of others for their dispersal. For example, the spores of *Streptomyces* can attach to the flagella of motile bacteria [18], while the coccoid cells of *Staphylococcus* attach directly to the cell body of swimming bacteria and can be passively translocated [42].

In our experiments, spherical cells of *C. pinensis* exhibited hitchhiking behaviour but unlike *Streptomyces* spores, this interaction was independent of carrier mobility and presence of flagella. Notably, hitchhiking was observed on swarming plates but not on swimming plates indicating that surface conditions influence this interaction (Figure 3) [43, 44]. In addition, gliding, sliding, and swarming are classified as crowd movements, whereas swimming is an individual behaviour [17, 45]. Crowd movements are an energy-saving, protective and efficient translocation mechanism that optimizes the search for resources in a nutrient-heterogeneous environment such as soil [37, 44, 46].

Microscopy imaging further supports that hitchhiking in *C. pinensis* is transient rather than a stable attachment, distinguishing if from *Streptomyces* spores. As observed in (Figure 3), spherical cells loosely associated with *B. subtilis* minicells, possibly via hydrophobic interactions. Notably, some spherical cells had already started elongating and losing their spherical morphology (Figure 3), but whether this promotes hitchhiking remains unclear.

In dense bacterial communities, population often align in liquid-crystal-like arrangements, promoting collective movement [47]. Consistent with this, *C. pinensis* spherical cells moved with *B. subtilis* rather than dispersing independently, despite being capable of surfactin cheating. Cheating on public goods, such as surfactin, is a well-known strategy in cooperative communities [39, 48]. However, *B. subtilis* employ regulatory mechanisms to mitigate cheater exploitation [39]. Thus, while hitchhiking behaviour was observed, it remains unclear whether *C. pinensis* actively engages in this process or undergoes passive translocation driven by *B. subtilis* crowd movements. Resolving this distinction is essential for understanding bacterial dispersal mechanisms and plant root colonization.

Each piece of knowledge contributes to unravelling and understanding the dynamic interaction within the host-microbiota. In addition, the question of the critical ratio of mobile and immobile bacteria that still allows community mobility should be answered. Considering that some bacteria are motile at low and others at high cell density, the question arises whether this contributes to the community remaining motile under different conditions. It would be also interesting to investigate the extent to which hitchhiking behaviour influences this movement, especially in the presence of plant roots. Some of these questions could potentially be explored in future studies using transparent soil and light-sheet fluorescence microscopy.

## Conclusion

Climate change and pollution from conventional agriculture degrade resource quality and put additional stress on plants, weakening their resilience. Preserving the natural occurring plant-protective microbes in the plant microbiota can improve plant resilience and immunity. To understand the complex interactions between plants and their microbiota and within the microbiota, the key members must first be identified and then their interaction mechanism uncovered.

One such key member is *C. pinensis*, known for its plant-health-promoting characteristics, demonstrated significant morphological plasticity transitioning from a long filamentous to a small spherical cell shape, accompanied by distinct transcriptional changes. Despite this transition, the spherical cells do not exhibit dormancy-like traits such as structural differentiation, increased resistance, or arrested growth but remain reproductive. In addition, the transition into small spherical cells enables flagella-independent translocation by motile bacteria.

Further work is needed to uncover the triggering and control mechanisms of morphological and metabolic plasticity and their role in dynamic and complex interactions. In addition, it is crucial to investigate the extent to which translocation, whether by hitchhiking or crowd movement, contributes to colonization and distribution within plant tissue. Deciphering these mechanisms enables harnessing the power of microbial communities to promote plant health and foster resilient agriculture in an increasingly challenging environment.

## Acknowledgements

Cryo-EM data were collected at the Netherlands Centre for Electron Nanoscopy (NeCEN) at Leiden University, with technical support from Dr. Wen Yang, Dr. Willem Noteborn, and Dr. Birgit Luef. This work benefited from access to NeCEN, which was funded in part by the Netherlands Electron Microscopy Infrastructure (NEMI), project number 184.034.014 of the National Roadmap for Large-Scale Research Infrastructure of the Dutch Research Council (NWO). J.L. was supported by the OCENW.GROOT.2019.063 and Building Blocks of Life 737.016.00 grants from the Netherlands Organization for Scientific Research (NWO), both awarded to A.B.

## Author Contributions

J.L. designed and conducted the study, performed most experiments, and wrote the manuscript draft. L.Z. contributed to laboratory work and provided input to the Methods section. D.U. supported the interpretation of transcriptomic data. F.R. assisted with statistical analysis and data processing. A.B. and G.P.v.W. supervised the project and provided critical feedback on the manuscript. All authors reviewed and approved the final version.

## Author Information

The authors declare no competing financial interests.

## Materials and Methods

### Strains and culturing conditions

*C. pinensis* 94 was obtained from culture collection of Carrion lab [7] and grown overnight from a 40% glycerol stock in 0.1x TSB at 25 °C and with agitation (250 rpm). After 16 h of growth, 100 µl of overnight culture was diluted into 50 ml 0.1x TSB and grown for an additional 20 h under the same conditions for obtaining long filamentous cells, and 40 h for small spherical cells (< 1 µm). Cells were harvested by centrifugation (30 min; 8000g; 4 °C). Subsequently, cell morphology was checked by light-microscopy (Axion Imager M2; Zeiss) before proceeding with any experiments. All *Bacillus subtilis* strains used in this study (undomesticated *Bacillus subtilis* (NCIB3610); *B. subtilis* DK605 ( Δ*mind\::TnYLB*); DS1677 (Δ*hag*); DS222 (Δ*motAB*); DK1484 (Δ*espH srfAA::mls*) were cultivated as described before [18]. *B. subtilis* mini-cell strain (DK605; Δ*mind::TnYLB*) was grown and harvested as previously described [18].

Prior to the experiments, the cell morphology of the cultures was assessed microscopically; in the following, the 20 and 40 h growth cultures are referred to as filamentous cells and spherical cells, unless otherwise stated. The experiments described below were carried out in triplicates (N = 3) unless otherwise stated.

### Quorum sensing assay

Cells of *C. pinensis* after 20 h (OD_600_= 0.2) and 40 h (OD OD_600_= 0.5) growth in 0.1x TSB (25 °C; 250 rpm) were harvested by centrifugation (8000x g; 30 min; 4 °C). Each sample was adjusted to an OD_600_ of 0.2 and 0.5 with fresh medium (0.1x TSB) and aliquoted with 1 ml. All sample aliquots were then centrifuged (8000x g; 30 min; 4 °C), the supernatant discarded, and the cell pellets used for the next step.

Preparation of media and culture supernatant for the identification of quorum-sensing signalling molecules. The supernatant of a 20 and 40 h *C. pinensis* culture was extracted by centrifugation (8000x g; 30 min; 4 °C) and filtration (0.2 µm). These media are referred to below as ‘long-filamentous cell media’ (LM) and ‘small spherical cell media’ (SM). Fresh TSB were diluted 1:10 with 40 h media and are referred to as ‘SM+T’ in the following. In addition, phosphate-buffered saline (PBS; pH 7) and fresh 0.1x TSB medium (FM) were used as reference media for the experiments. The experimental setup is illustrated in Figure S2. Aliquots of each sample were resuspended in 1 ml of the respective prepared medium and transferred to 96-well plates (MegaBlock®; 2.2 ml; Sarstedt), which were subsequently incubated at 25 °C and 250 rpm.

After 3 and 6 h, samples were taken from all wells, treated with Syto9 (1 µM) and fixed with 4% paraformaldehyde. All samples were imaged with an upright fluorescence microscope (Zeiss Axio Imager M2 with AxioCam MRc 5), and the cell length data was determined with ImageJ [49] using the AutoTreshold function. The resulting data were visualized graphically and *p*-values were obtained from a mixed effects regression model, using the logarithm of the cell length to obtain approximate conditional normality and accounting for dependency by including a random effect for “replicate". Mixed models were fit using the lme4 package, and *p*-values were obtained using the lmerTest and emmeans packages lme4 [50], lmerTest [51], emmeans [52].

### Motility assay

#### Hitchhiking motility assay

The motility test was performed on 0.1x TSB agar plates (ø 9 cm) with 0.5% agar for swarming plates and 0.27 – 0.3% agar for swimming plate assays. First, a new culture of *B. subtilis* and its mutants was prepared from an overnight LB culture and incubated in LB (37 °C; 200 rpm) until an OD_600_ of 0.5 was reached. Subsequently, *C. pinensis* cells were harvested from plate (0.1x TSA; 25 °C; > 1-week incubation) using a sterile loop and resuspended in 200 µl 0.1x TSB. To determine the hitchhiking behaviour, 3 µl of *B. subtilis* cells were transferred to the plate and 3 µl of *C. pinensis* cell suspension was pipetted directly onto the desired inoculation site (either directly on the *B. subtilis* inoculation site or onto a separate site on the plate). For the hitchhiking assay on swimming plates (0.27–0.3% agar; LP0011; Oxoid), a sterile loop was punctured into each inoculation site after inoculation to promote swimming behaviour inside the agar. The plates were incubated at 25 °C for up to 5 days and imaged on a light box. Experiments were performed in triplicate using Bacto^TM^ Agar (BD) and Agar No. 1 (LP0011; Oxoid). For comparison and control purposes, each condition was also tested on each strain individually.

#### Surfactin cheating assay

Surfactin cheating assays were performed on 0.1x TSA (0.5% agar) with surfactin from *B. subtilis* (Sigma-Aldrich; St. Luis; Missouri; USA). For the incorporation of surfactant into the medium, 10 ml medium was liquefied by heating in a microwave and 10 µl surfactin (stock: 10 g/L) added after cooling down to room temperature. For the cheating assay in combination with additional potassium, 70 µl of K2HPO4 (1 M) was added to the liquid medium. The medium was then poured into a petri dish (ø 9 cm) and solidified. The topical treatment of media plates was performed by pipetting 10 µl of surfactin (10 g/L) onto the centre of the plate and left under the hood until the surfactin had completely diffused into the agar. The procedure was repeated with 1 g/L surfactin solution. All plates were inoculated with a single colony of *C. pinensis*, taken with a sterile tooth pick from a starting plate (0.1x TSA; 1.5% agar). The plates were incubated at 25 °C for 3 days and imaged on a light box. The experiments were carried out in triplicate. For comparison, media without surfactin and additional potassium were inoculated and treated in the same way.

### Fluorescence microscopy

Fluorescence microscopy was performed to evaluate and confirm the successful integration of GFP into the genome of *C. pinensis*. For this purpose, a colony was sampled from a plate, suspended in 3–5 µl PBS on a glass slide, and covered with a coverslip. Microscopy and imaging were carried out using a Zeiss Axio Imager M2 equipped with an AxioCam MRc 5 camera. GFP fluorescence was imaged (Ex 488 nm; Em 510 nm) and analysed in Zeiss Zen software.

### Cryo-electron microscopy

Cell pellet obtained from *B. subtilis* mini-cell strain (DK605; Δ *mind::TnYLB*) as described above was resuspended in 12 µl LB. *C. pinensis* spherical cell morphology were harvested with a sterile loop from a 2-week-old 0.1x TSB plate (1.5% agar) and transferred to 200 µl 0.1x TSB medium. From the *C. pinensis* cell suspension, 2 µl were added to the 12 µl *B. subtilis* mini-cells and a 10-nm colloidal gold solution treated with protein A (Cell Microscopy Core, Utrecht University, Utrecht, The Netherlands) was added in a 1/10 dilution. The mixture was gently mixed by pipetting and incubated for 30 minutes at RT. Subsequently, 3 µl of sample were applied to glow-discharged 200-mesh R2/2 copper grid (Quantifoil Micro tools), pre-blotted for 30 sec. and then blotted for 1 sec. in a chamber with 90% humidity and at 20 °C. The grids were plunge frozen in liquid ethane using an automated Leica EM GP system (Leica Microsystems) and then stored in liquid nitrogen. Imaging of the grids was performed with a 120 kV Talos L120C cryo-electron microscope (Thermo Fisher Scientific) at the Netherlands Centre for Electron Nanoscopy (NECEN).

### Cryo-electron tomography

The *C. pinensis* cells were prepared as described above and samples were taken after 20 h and 40 h incubation. The samples were then centrifuged at 3000x g for 5 min. and the cell pellet were resuspended in 20 µl. In addition, a 10-nm solution of colloidal gold treated with protein A (Cell Microscopy Core, Utrecht University, Utrecht, Netherlands) was added in a 1/10 dilution and carefully mixed by pipetting. For vitrification, 3 µl of sample was applied to a glow-discharged grid and plunge frozen as described above. Imaging data were acquired using a TITAN Krios microscope (TFS), equipped with a Gatan K3 Summit direct electron detector and a GATN GIF Quantum energy filter with a slit width of 20 eV. Images were acquired at a nominal magnification of 26,000, which corresponds to a pixel size of 3.28 Å. Tilt series were collected using SerialEM with a bidirectional dose-symmetric tilt scheme (−60° to 60; starting from 0°) with a 2° increment. The defocus was set between -8 and -10 µm, and the cumulative exposure per tilt series was 100 e-/A2. Alignment of the tilt series based on bead tracking and drift correction was performed with IMOD [53] and CTFplotter was used to determine and correct the contrast transfer function [54].

## Supplementary Figures

**Figure S1.**
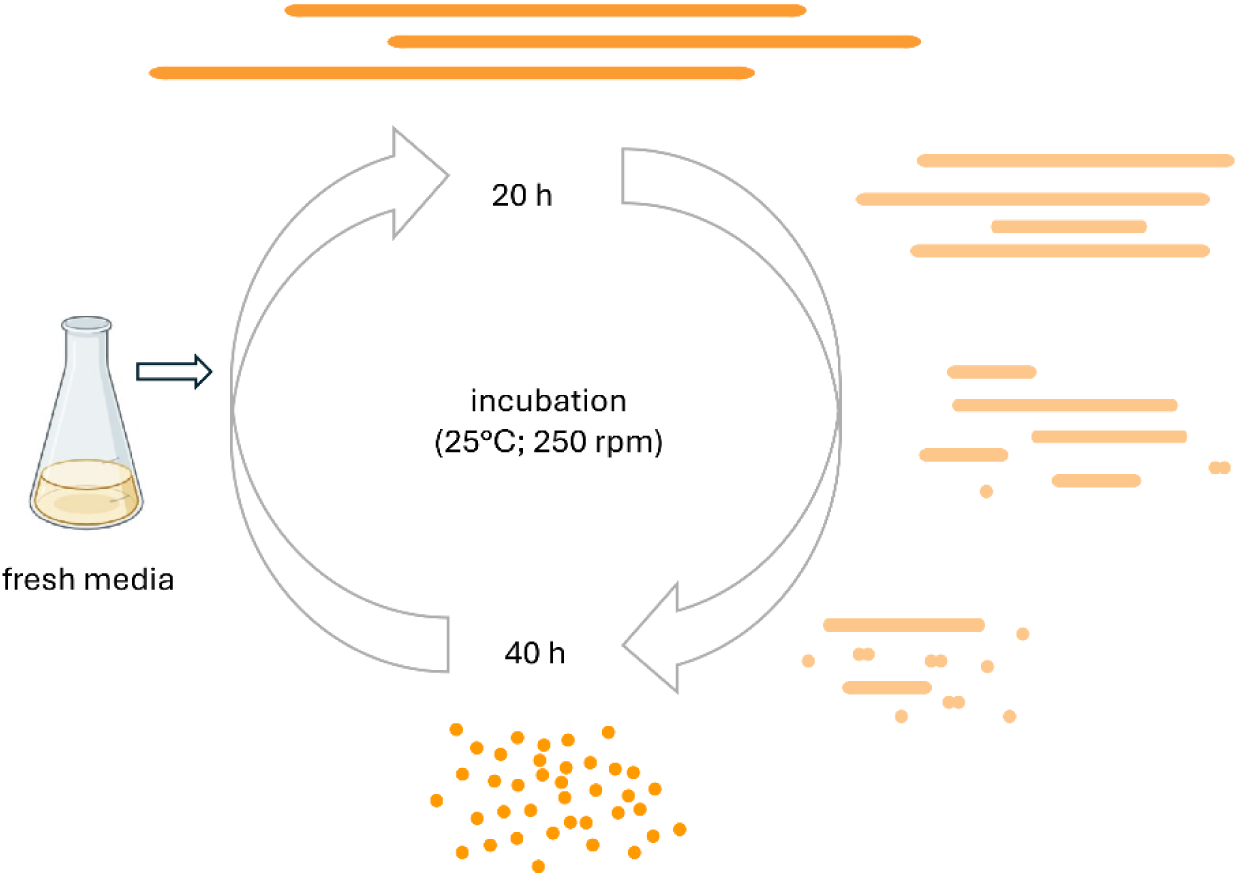
Schematic representation of the reversible morphological cycle of *C. pinensis* during growth in liquid culture (25 °C, 250 rpm). The culture transitions from predominantly spherical cells to filamentous forms within 20 h and returns to spherical morphology after 40 h. Intermediate stages are shown in lighter colour to reflect the unresolved timing of the transition. Attempts to synchronize the cell cycle and capture these intermediate forms were unsuccessful, preventing accurate determination of timing and further analysis.

**Figure S2.**
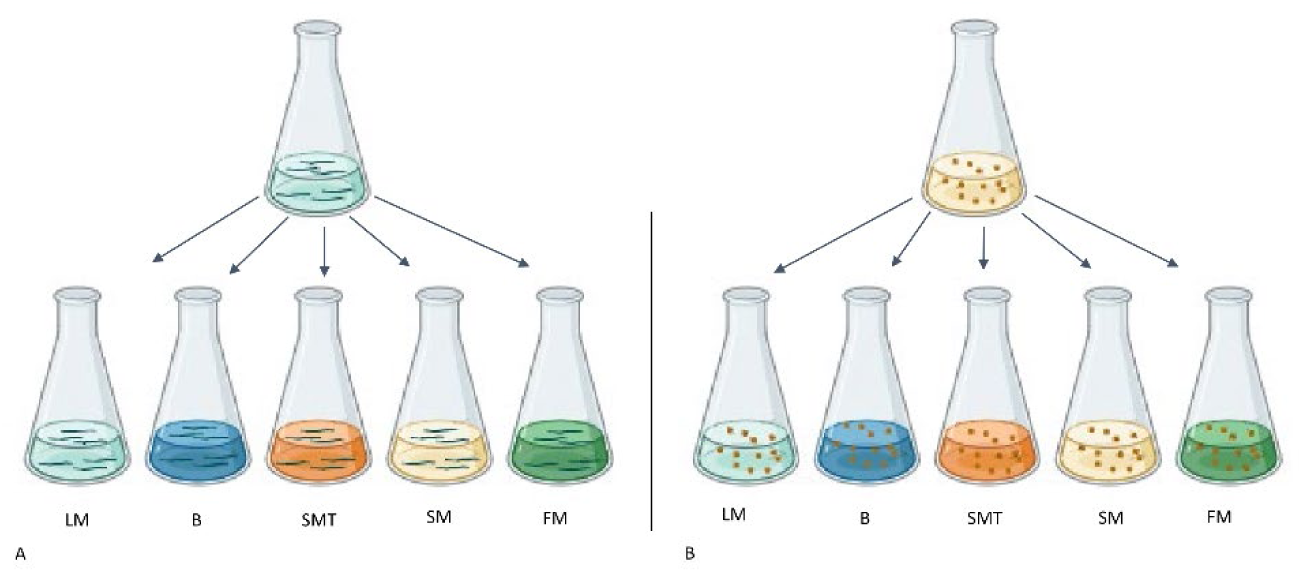
Experimental setup of the quorum sensing assay. Long filamentous cells (LC) of a *C. pinensis* culture after 20 h incubation (25 °C; 250 rpm) and small spherical cells (SC) after 40 h of incubation in 0.1x TSB were harvested. Following respective supernatant of the 20 h (LM) and the 40 h (SM) culture was sterile filtrated and used in the setup, whereby one part of the SM supernatant was supplemented with full TSB media in a 1:10 dilution (SMT). Phosphate-buffered saline (B) and fresh 0.1x TSB medium (FM) were included as reference medium. The initial cell density for both cell morphologies were setup to an OD_600_ of 0.2 and at a higher OD_600_ of 0.5. The experiment was conducted in triplicates.

**Figure S3.**
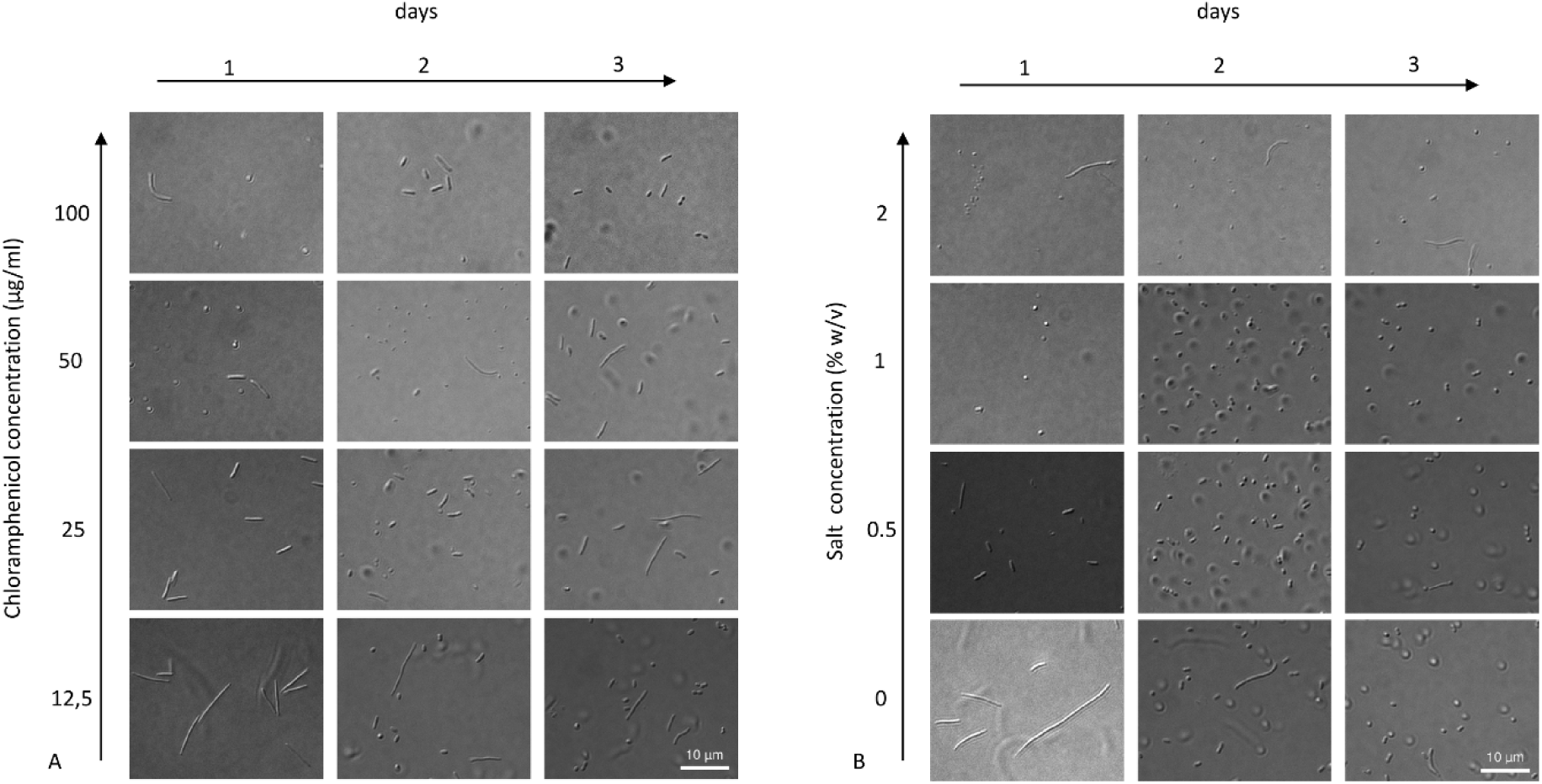
Stress reaction test of *C. pinensis* (N= 3) in 0.1x TSB with increasing chloramphenicol (µg/ml) and salt concentration (% w/v). To investigate the extent to which the formation of small spherical cells is a stress response of *C. pinensis*, cells were first harvested at the long filamentous cell stage after 20 hours of incubation and then exposed to chloramphenicol concentrations ranging from 12.5 µg/ml to 100 µg/ml (A). Chloramphenicol is an antibiotic that binds to the 50 ribosomal subunit and thus impairs protein biosynthesis. In addition, the cells were exposed to salt concentrations of 0.5% to 2% (B), which disrupt the integrity of the cell wall. A culture in 0.1x TSB without additional salt was used as a reference sample (B). Cells were imaged every 20 h for 3 days. Scale bar 10 µm.

**Figure S4.**
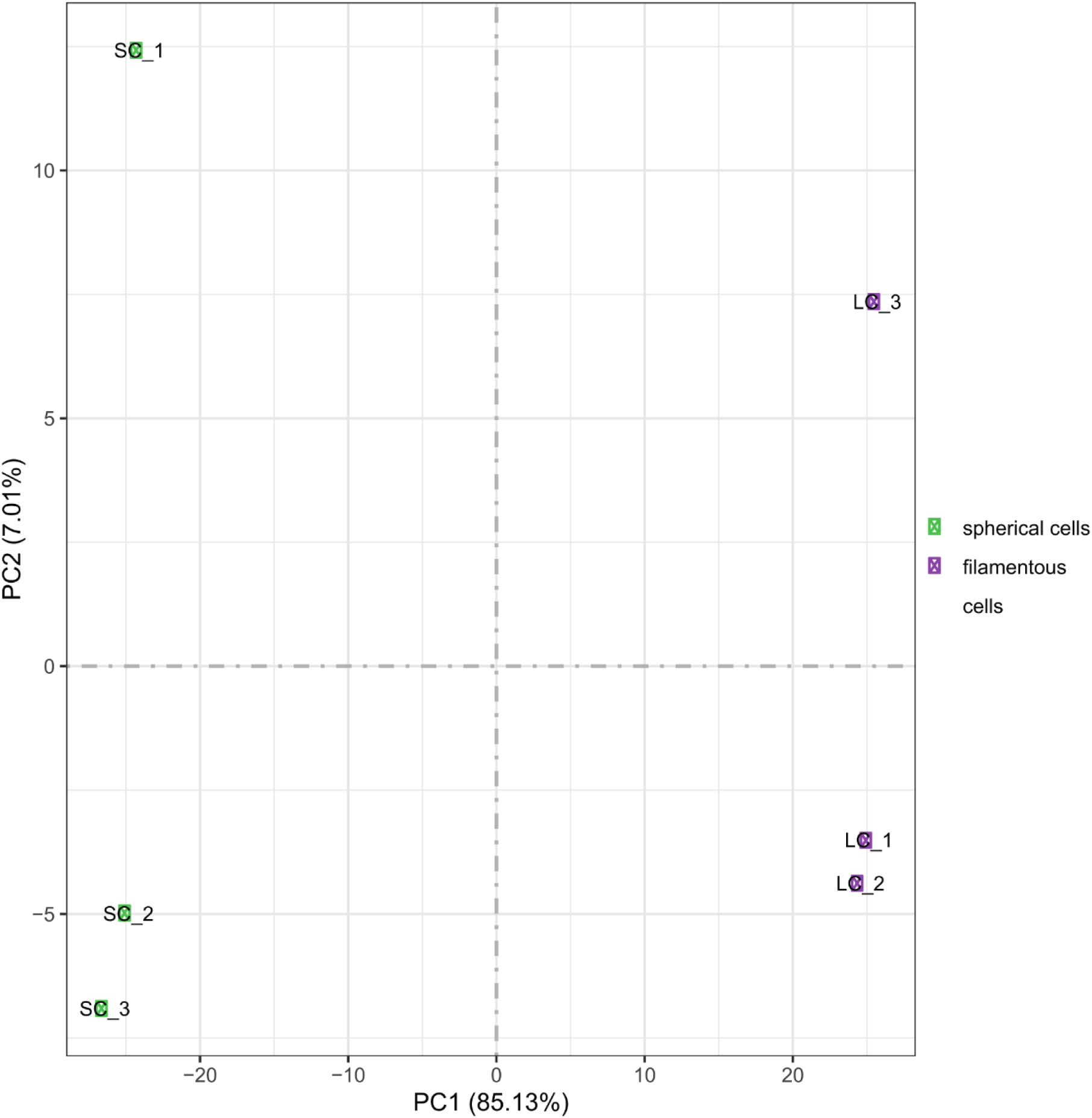
Principal component analysis (PCA) of transcriptomic profiles from *C. pinensis* cell morphologies. Samples cluster according to morphology, with PC1 explaining 85.13% of variance, separating spherical cell (SC) from filamentous cells (LC), indicating distinct transcriptional states.

**Figure S5.**
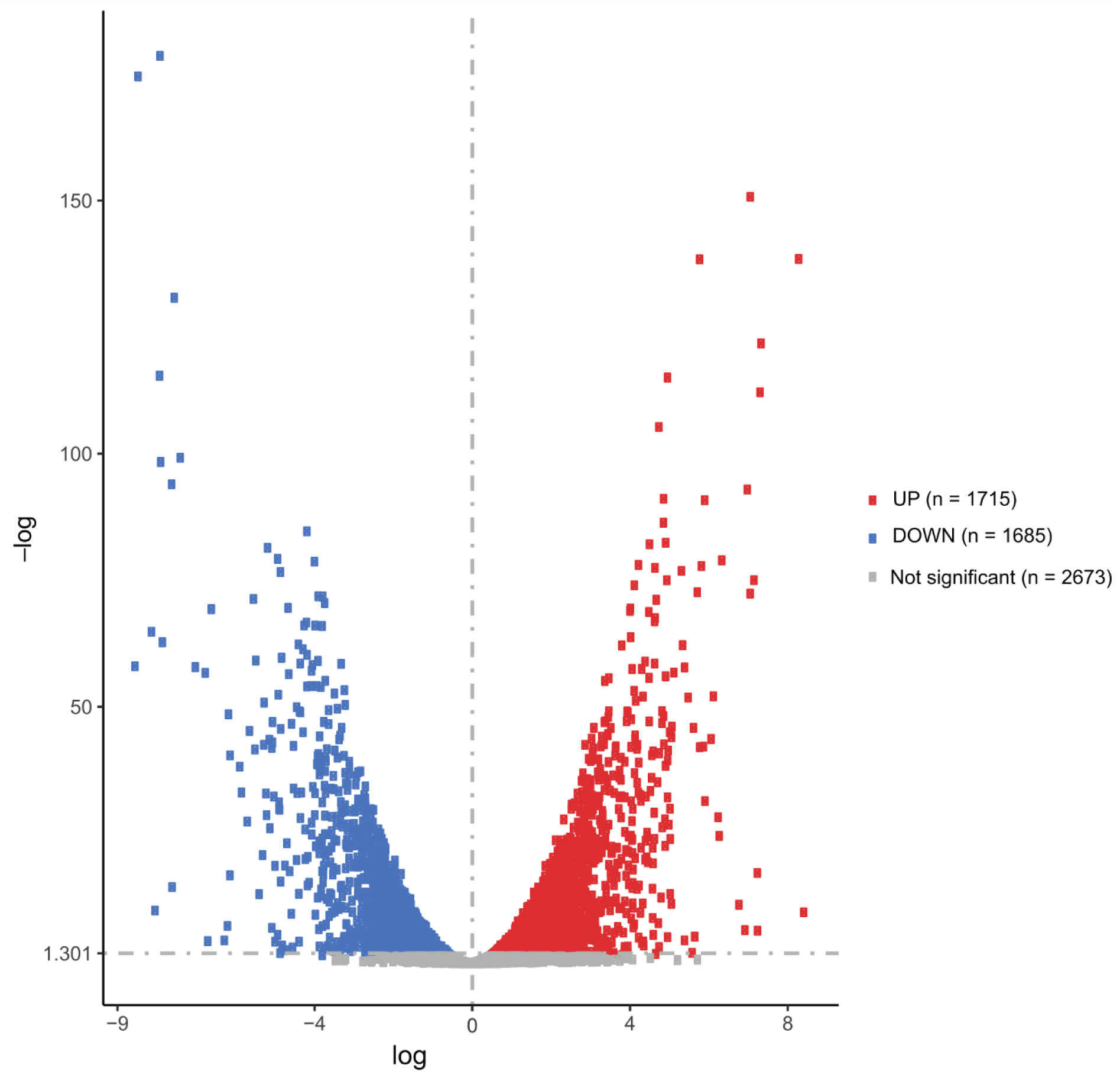
Volcano plot of differential gene expression (DEG) between spherical and filamentous *C. pinensis* cells. Genes with a fold change of 2 (FC ≥ 2) and an adjusted *p*-value (padj) of 0.05 are considered significantly regulated: red = upregulated (n = 1715), green = downregulated (n = 1685), blue = non-significant (n = 2673).

**Figure S6.**
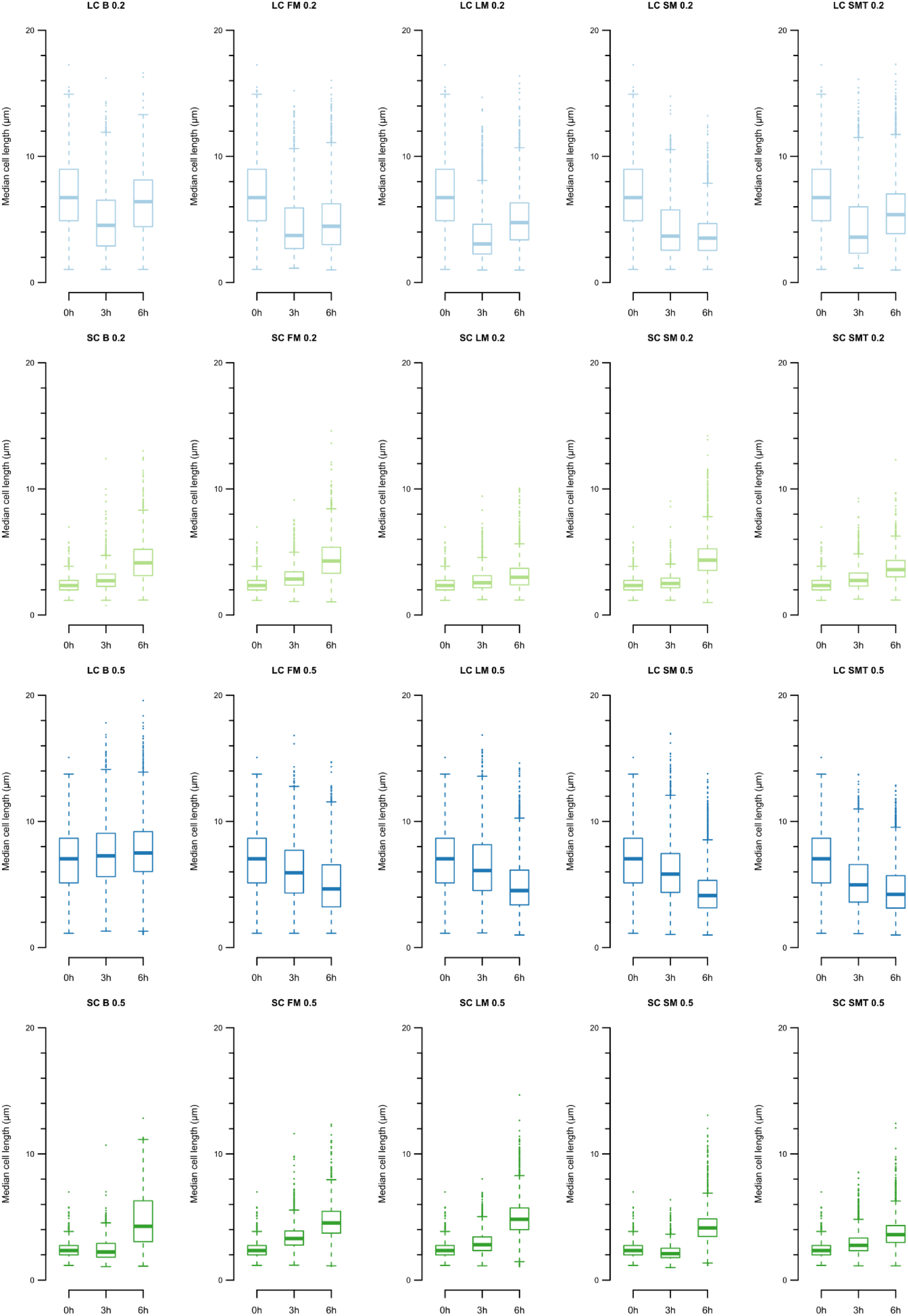
Comparison of *C. pinensis* morphological change over time under different growth conditions. (A - E; K - O) Change from an initial long filamentous cell shape (LC) with an initial cell density of OD_600_ = 0.2 (A-E) and 0.5 (K - O). (F – J; P – T) As well the transition from a small spherical cell shape (SC) with an initial cell density of OD_600_ = 0.2 (F – J) and 0.5 (P – T). Following conditions were used: buffer (B); fresh medium (FM); sterile filtrated supernatant of 20 h growth medium (LM); sterile filtrated supernatant of 40 h growth medium (SM) and sterile filtrated supernatant of 40 h growth medium supplemented with full medium (SMT). Cell lengths (µm) were measured after 3 and 6 hours after inoculation. Confidence interval 95% and N=3.

## Supplementary Methods

### Physical stressor assay

To measure the differences in viability of the filamentous and spherical cell morphologies when exposed to physical and chemical stresses, we first harvested the cells from liquid culture (50 ml) by centrifugation for 10 min. (filamentous cells) and 30 min. (spherical cells) at 8,000x g and at 4 °C. All samples were adjusted to an OD_600_ of 0.2 and stored on ice until the various treatments outlined below. Subsequently, 5 µl of each sample was drop plated in triplicates on 0.1x TSA plates (1.5% agar) and incubated at 25 °C. The plates were imaged on a light box after 2 days.

#### Treatment with ultrasonic

For this treatment, 1 ml of each sample was transferred to a 15 ml Falcon tube and treated in an ultrasonic bath (ultrasonic cleaner; CDS-100; AC 220 240 V 50/60 HZ; 42,000 hz) for 2 or 5 min. and then placed back on ice.

#### Heat treatment (60 °C)

1 ml of each sample was transferred to a Falcon tube (15 ml) and placed in a 60 °C pre-warmed water bath (Type: W 200; 220 V; 5.5 A; 50 Hz; 1200 W; Memmert GmbH + Co KG) for 2 or 5 min. and then placed back on ice.

#### Treatment with ultraviolet light (UV)

UV-light treatment of each sample was performed as described before [55] in a biosafety cabinet (BSC-700II-I; HMC Europe) with a UV-light source (ZW15S19W-Z436; Cnlight; Shelley). From each sample, 5 ml of the culture was transferred to a sterile petri dish (Ø 9 cm) and placed without a lid at a distance of 50 cm from the UV-light source. The exposure time was 5 or 10 min., and after treatment the samples were cooled at room temperature before plating.

#### Desiccation test

Filamentous and spherical cells were plated on 0.1x TSA and incubated at 25 °C for 8 weeks until the agar had completely dried out. Subsequently, 1 ml of 0.1x TSB and a sterile loop was used to scrape the cells from dried agar plates. From the resulting cell suspension, 100 µl was used to inoculate and spread onto new 0.1x TSA media plates. The plates were incubated at 25 °C and imaged after 2 weeks.

### Chemical stressor assay

The experiments to test the resistance behaviour of the two cell morphologies were carried out on 96-well plates and with biological and technical triplicates. The cells were harvested as described above and the OD_600_ of each sample was set to 0.5 – 0.6. Each well was prefilled with 180 µl 0.1x TSB and 20 µl of the cell cultures were added. The measurement was performed in an automated Spark® multimode microplate reader (Tecan, Switzerland) with the following parameters: Temperature 25 °C; measuring time 22 - 24 h; amplitude 2 mm; frequency 810 rpm; duration 10 s; linear duration 5 s; wavelength 600 nm; 10 flashes; settle time 5 ms and an interval of 30 minutes. For the evaluation and quantification of the growth parameters under the different chemical growth conditions, the doubling time, growth rate and carrying capacity were computed in R (version 4.3.2) using the packages growthcurver (version: 0.3.1) and lme4 (version 1.1-35.1) and the RStudio IDE.

#### Pondus Hydrogenii (pH) test

The pH test was carried out with 0.1x TSB medium (pH 7.2), which was adjusted to pH 5 and pH 8 with HCl and NaOH. Untreated 0.1x TSB (pH 7.2) was used as a positive and growth control.

#### Sodium dodecyl sulfate (SDS)

Sterile filtrated SDS (dissolved in 0.1x TSB) was added to the medium (0.1x TSB) at a final concentration of 0.02%; 0.2%; and 2% (w/v).

#### Ethanol test

Ethanol was added to the medium (0.1x TSB) at a final concentration of 0.02%; 0.2%; and 2% (w/v).

### Stress-reaction test

Cultures of *C. pinensis* (25 °C; 250 rpm; 20 h; OD_600_ = 0.1) were centrifuged (1500x g; 30 min; 4 °C) to obtain cell pellets, which were resuspended in the following media:

Liquid media with different chloramphenicol concentrations were prepared with a chloramphenicol stock solution (25 mg/ml) and diluted with 0.1x TSB to final concentration of 100, 50, 25 and 12.5 µg/ml chloramphenicol. Media with different NaCl concentrations (0.5 – 2%) were prepared from a 10% (w/v) stock solution in 0.1x TSB. In addition, 0.1x TSB was used as the medium for the oxygen stress test. Subsequently, 1 ml of each sample was transferred to a 96-DeepWell^TM^ plate and incubated (25 °C; 250 rpm; 3 days). To prevent evaporation of the medium, the plates were sealed with a sealing film, whereby the wells of the oxygen stress test were sealed with 3 layers of sealing film. To prevent additional oxygen from entering the oxygen stress test, the sealing film was only cut open for one row of wells for each examination. Every 20 h, 10 µl was taken from each sample and the cell morphology was examined using a light microscope (Axio Imager M2; Zeiss). All tests were performed in triplicates.

### Transcriptomics and data analysis

*C. pinensis* was cultured in 50 ml 0.1x TSB (25 °C; 250 rpm), and cells were harvested by centrifugation (1500x g; 15 min; 4 °C) after 20 h (filamentous cells; OD_600_: 0.16 ± 0.01) and 40 h (spherical cells; OD_600_: 0.9 ± 0.1) of growth. Due to the low density, the cell pellets of 3x 50 ml *C. pinensis* 20 h culture were pooled to obtain a sufficient number of cells for RNA extraction. From the 40 h *C. pinensis* culture, 10 ml were used for cell harvesting. The cell pellets were resuspended in 1 ml RNAprotect®, incubated for 5 minutes at room temperature (RT) and then washed with 0.1x TSB (centrifugation: 5000x g; 10 min; RT). Total RNA was extracted and purified using the RNeasy® Mini Kit (Qiagen; Hilden; Germany) according to the manufacturer’s instructions. The removal of rRNA, library preparation, Illumina sequencing and subsequent quality control of the data were performed by Novogene Europe (Cambridge, UK). The RNA-sequence data was analysed using a standardized pipeline. The raw data was processed to remove adapter sequences and low-quality reads. The sequences were then aligned to the reference genome *C. pinensis* (accession no. LR632929 from the study [7]) annotated by Prokka using Bowtie2. Gene expression levels were quantified using FeatureCounts and differential expression analysis was performed using DESeq2 (1.20.0). Genes with an adjusted *p*-value below 0.05, as determined by DESeq2, were considered differentially expressed. Gene Ontology and KEGG pathway enrichment analyses were performed with clusterProfiler (v3.8.1).

### Expression of green fluorescent protein (GFP)

Native *gap* promoter was amplified from genomic DNA of *C. pinensis* using primer pair Pgap_chi_F (5’- AGGGAATTCCGGACCGGTACCCCACAGGTCGCCCATAAATAA-3’) and Pgap_chi_R

(5’- TTCTTCTCCTTTACTCATTTTACACTGTAGTTTGTAGATGAAAAATTGAG-3’), gene gfpmut3 was amplified from synthesized DNA fragment harbouring gfpmut3 [56] using primers gfpmut3_F (5’- ACAGTGTAAAATGAGTAAAGGAGAAGAACTTTTCACT-3’) and Chitin_GFP_R

(5’- TGCATGCCTGCAGGTCGACTCTAGATATTTGTCCTACTCAGGAGAGCGTTC-3’), the aforementioned PCR products were purified and cloned into KpnI and XbaI digested pCP23 [57] via Gibson assembly. The generated construct pGWS1802, which expresses GFPmut3 under the control of the *gap* promoter, was introduced into *C. pinensis* by electroporation as described previously [56]. The successful introduction of pGWS1802 was validated by colony PCR on transformants which were able to grow on CYE agar containing tetracycline (100 µg/mL), using primers pair gfpmut3_F and Chitin_GFP_R.

